# Inferring the stabilization effects of SARS-CoV-2 variants on the binding with ACE2 receptor

**DOI:** 10.1101/2021.04.18.440345

**Authors:** Mattia Miotto, Lorenzo Di Rienzo, Giorgio Gosti, Leonardo Bo’, Giacomo Parisi, Roberta Piacentini, Alberto Boffi, Giancarlo Ruocco, Edoardo Milanetti

## Abstract

With the progression of the SARS-CoV-2 (severe acute respiratory syndrome coronavirus 2) pandemic, several variants of the virus are emerging with mutations distributed all over the viral sequence. While most of them are expected to have little to no effects at the phenotype level, some of these variants presenting specific mutations on the Spike protein are rapidly spreading, making urgent the need of characterizing their effects on phenotype features like contagiousness and antigenicity. With this aim, we performed extensive molecular dynamics simulations on a selected set of possible Spike variants in order to assess the stabilizing effect of particular amino acid substitutions, with a special focus on the mutations that are both characteristic of the top three most worrying variants at the moment, i.e the English, South African and Amazonian ones, and that occur at the molecular interface between SARS-CoV-2 Spike protein and its human ACE2 receptor. We characterize these variants’ effect in terms of (i) residues mobility, (ii) compactness, studying the network of interactions at the interface, and (iii) variation of shape complementarity via expanding the molecular surfaces in the Zernike basis. Overall, our analyses highlighted greater stability of the three variant complexes with respect to both the wild type and two negative control systems, especially for the English and Amazonian variants. In addition, in the three variants, we investigate the effects a not-yet observed mutation in position 501 could provoke on complex stability. We found that a phenylalanine mutation behaves similarly to the English variant and may cooperate in further increasing the stability of the South African one, hinting at the need for careful surveillance for the emergence of such kind of mutations in the population. Ultimately, we show that the observables we propose describe key features for the stability of the ACE2-spike complex and can help to monitor further possible spike variants.

## I. INTRODUCTION

The severe acute respiratory syndrome coronavirus 2 (SARS-CoV-2) infection was firstly observed in late 2019 [1, 2]. In the subsequent months of epidemic spreading, many SARS-CoV-2 variants, viral sequences characterized by at least one mutation with respect to the original one, have been detected worldwide [3].

In coronaviruses, mutations naturally occur during viral replication, and despite the fact that coronaviruses encode for an enzyme that corrects the errors made some of this mutation are preserved forming new variants. As for all biological systems, the action of natural selection eventually tends to fix in the genome the mutations characteristic of those variants that present an increase of the fitness, and it has been registered that in these months the rate of emergence of new SARS-CoV-2 variants is about two variants per month [4, 5].

This rapid proliferation of variants poses a further threat for the community as the virus can acquire different phenotypes.

For instance, the main mutation of the B line, involving the amino acid substitution D614G in Spike, is established since March 2020 and it is now largely dominant in patients [6–8]. This mutation would allow the receptor-binding domain (RBD) of Spike to assume a conformation more suitable to bind Angiotensin-converting enzyme 2 (ACE2), and it could be responsible for the increased viral action [9, 10].

More recently, on December 14, 2020, authorities in the United Kingdom of Great Britain and Northern Ireland reported to World Health Organization that a new variant SARS-CoV-2, B.1.1.7, commonly known as the English variant, has been identified via viral genomic sequencing [11, 12]. The variant is defined by the presence of a range of 14 mutations involving amino acid modifications and three deletions, including the spike D614G mutation. Even if investigations are underway to determine whether this variant is associated with any changes in antibody response or vaccine efficacy, it seems to be characterized by an increased transmissibility [13] and lethality [14], also because it is spreading with very high speed all over the world.

Among the mutations affecting the spike protein, like 69-70del or P681H, mutation N501Y is located in the region that directly contacts the ACE2 receptor. Therefore, it is possible that this mutation could have a direct effect on the binding affinity between the two proteins [15].

Furthermore, the importance of this amino acid substitution is confirmed by its presence in two other rapidly spreading variants, B.1.351 [16] and P.1 [17] (commonly referred to as South African variant and Amazonian variant, respectively).

These two variants both include other amino acid substitutions in the spike binding site, making clear that this region is under severe evolutionary pressure. Indeed, selectivity and the affinity of the spike toward its main receptor, ACE2, remain the crucial factors determining SARS-CoV-2 contagiousness and virulence.

Even if, it has been demonstrated that SARS-CoV-2, similarly to other coronaviruses [18, 19], is able to bind to sialoglycan-based receptors in the N-terminal domain [20, 21].

From a molecular point of view, the spike protein of CoVs, protruding from the viral membrane, not only plays a crucial role as a fundamental structural protein, but it also is essential for the interaction between CoV systems and host cells [22]. Structurally, the spike protein is found in the trimeric complex, each chain composed of two sub-units: S1 and S2. The Receptor Binding Domain (RBD), located in the S1 domain, is responsible for viruses’ interaction with receptors on the host cell surface [23]. On the other hand, the S2 subunit is responsible for the fusion between the virus and host membrane, causing the viral genome to penetrate the host cell’s cytoplasm [24].

Interestingly, the interaction with ACE2 involves the C-terminal domain of SARS-CoV-2 spike protein, whose amino acid sequence is well conserved with respect to SARS-CoV homologous one [25]. Conversely, the N-terminal domain presents some insertions, and these additional surface regions could be used by the virus to bind other cell receptors, so constituting an additional cell entry mechanism [26].

Here, we perform extensive molecular dynamics simulation of different systems of the spike-receptor complex, involving mutations at the spike protein interface. In particular, we both consider single mutations belonging to the binding site of the spike protein for which the binding affinity experimental data is known, and we consider the variants of SARS-CoV-2 currently most widespread in the world. We analyze the mutations for which binding affinity is known in order to detect the dynamic-structural properties of the ACE2-spike complex that give the spike protein a greater propensity to interact with the molecular partner. Only by considering the RBD of the spike protein and the extra-membrane domain of the ACE2 receptor, we analyze the dynamic properties of the residues belonging to the two interfaces. Furthermore, an analysis based on graph theory was performed in order to investigate the stability of the contacts during molecular dynamics simulations. Finally, we analyze the shape complementarity of the two interfaces over time, using Zernike polynomials to characterize the shape of each portion of the molecular surface, establishing a complementarity value between each pair of surface [27]. Also, in this case, we investigate the stability of the geometrical matching between the two interfaces during the molecular dynamics simulation because stable binding to the host receptor plays a crucial role for virus entry mechanisms [28].

This study shows that the English variant and the South African variant have structural and dynamic properties similar to the N501F variant, which is experimentally known to be the one with the highest binding affinity with ACE2. This finding invites us to explore the possible cooperative effect of mutations in the binding region. Therefore, we substituted the F amino acid (Phenylalanine) at position 501 of the spike protein for both the Amazon variant and the South African variant. The results obtained from our fully computational approach show that the presence of the F amino acid in the Amazonian variant would worsen the binding affinity. On the other hand, the same mutation carried out in addition to the mutations of the South African variant would increase the stability of the spike-ACE2 complex, suggesting these mutations as a possible worrying variant.

## II. RESULTS AND DISCUSSION

### A. The role of spike mutations in the emerging variants

Although mutations are showing up all over the SARS-CoV-2 viral genome, those taking place on the spike protein are under intense scrutiny as they are expected to directly impact viral entrance in the host cells, thus on transmissibility and infectivity. The first notable mutation that has been observed involved residue 614 which changed from D to G [6]. This mutation even if localized in a region distant from the binding regions, rapidly fixed in the population suggesting an indirect effect on the phenotype. A recent computation study highlighted a conformational change driven by the mutation that favors ACE2 receptor binding, thus explaining the phenotypic advantage [10].

With the huge spread of the epidemics, other mutations accumulated over the SARS-CoV-2 genome. In particular, a variant of the virus with 6 mutated aminoacid and 2 deletions in the spike protein emerged in England in the past mounts. The English variant has the G614D mutation together with mutations/deletions belonging to spike protein: 69-70 HV deletion, 144Y deletion, N501Y, A570D, P681H, T716I, S982A, and D1118H. Specifically, mutation N501Y and deletions 69-70 attracted much attention since the mutation falls in the receptor-binding domain while the double deletion interests the N-terminal domain, which has been demonstrated to bind sialic acid-rich attachment factors [20, 21].

To begin with, to assess the variance invasion potential we looked at genomic data taken from infected individuals divided by geographical location, which is provided by the regularly updated public web site (**https://cov.lanl.gov**). In particular, analyses are based on 823,121 spike alignment sequences taken at different times and different geographical areas using the consistency and significance test presented by Korber *et al*. [29]. First, we compared sequences carrying the ’wild type’ residue in position 501 (i.e. N501) with the ones carrying the 501Y mutation.

Figure 1a displays the changes of relative frequencies for N501 and Y501 for two-time points separated by two weeks. The first time point represents all sequences up to the onset day (at least 15 sequences required with 3 least represented sequences), and the second time point includes all the sequences acquired at least two weeks after the onset date (at least 15 sequences required). One can see that in each country, the variant 501Y is rapidly spreading.

**FIG. 1:**
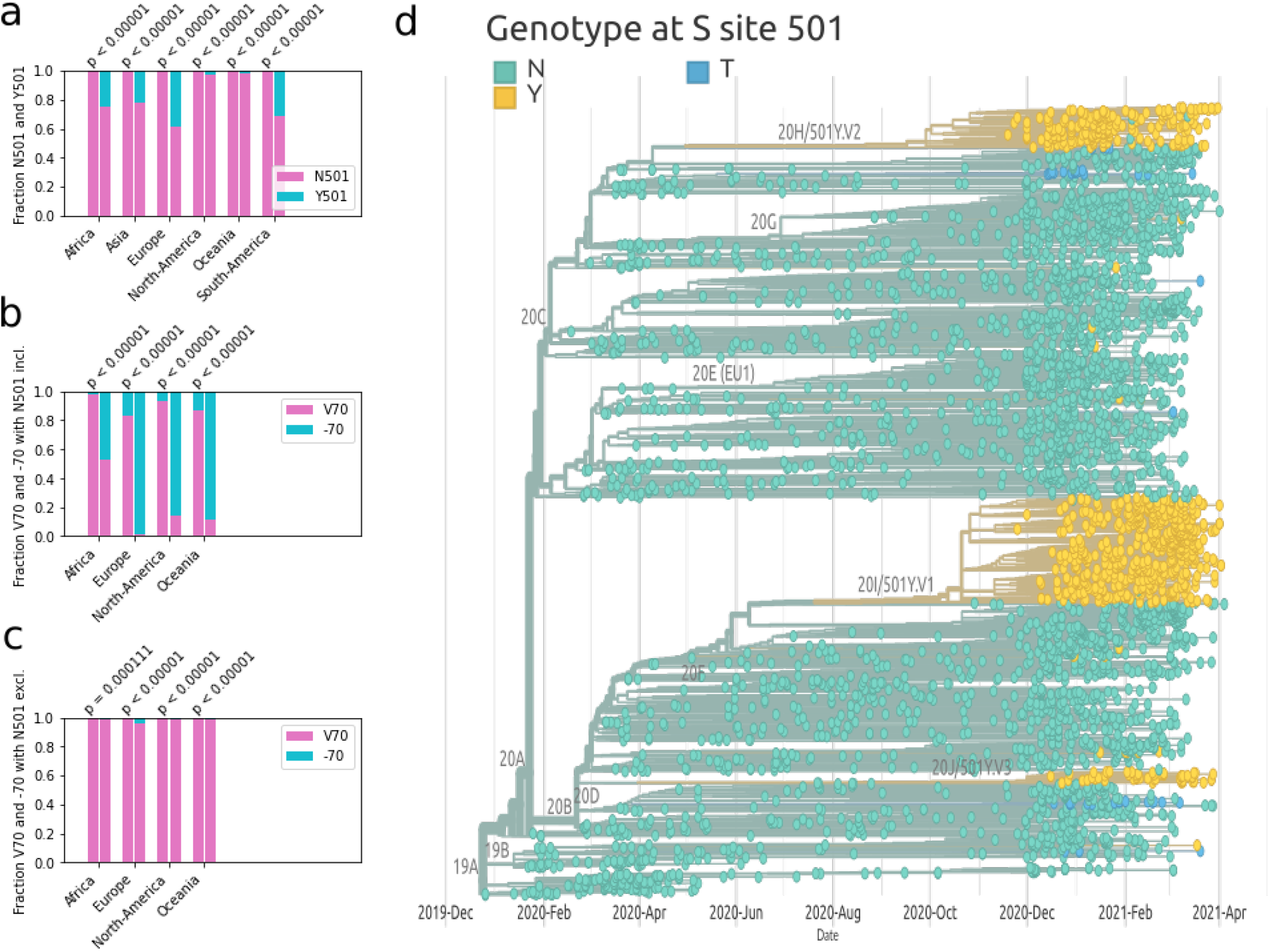
**a)**Fractions of sequenced viral samples having amino acid N in position 501 of the spike protein (orange) vs those having Y (blue) for two different time intervals and in different countries. **b)** Same as panel a), but comparing samples with residue V70 (orange) and samples for which deletion of residue 70 took place (blue). **c)** Same as in panel b) but considering only sequences where residue 501 is an N. **d)** Phylogenetic tree of the SARS-CoV-2 variants. Leaves are colored in cyan if the corresponding spike sequence has an N in position 501, in dark blue in case of a T, and in yellow for a Y.

The significance of the signal is obtained testing the null hypothesis that the mutation does not affect the variant fitness and that consequently, the relative frequency shift must randomly go with equal probability in each direction. The p-values confirm that in all continents we observe a systematic increase of the relative frequency of Y501. Next, we moved to consider sequences carrying the deletion V70 (taking place on the glycan-binding domain). Figures 1b-c show respectively the change in the relative frequency of the deletion at position 70 constraining respectively for the N501 and Y501 variant; this shows that the deletion of the 70th position increases in relative frequencies at a much greater rate in the Y501 variant. This suggests that deletion 70 is driven by the rapid spread of the English variant, and/or that the Y501 variant gives a stronger relative change in fitness to the deletion.

Finally, looking at all infection data, we compare the invasive potential of the different 501 mutations. The phylogenetic tree in Figure 1a was obtained using the publicly available interactive visualization platform Nextstrain [30] and it is based on the contribution of 4025 genomes sampled between December 2019 and April 2021, stored and elaborated in the Nextstation database and bioinformatics pipeline for phylodynamics analysis [30]. A full list of the sequence authors is available at **https://nextstrain.org/sars-cov-2**. Light green dots represent sampled genomes with N501 sequences, yellow dots Y501 and blue T501. One can see that the N501Y mutation independently emerged in different branches of the phylogenetic tree and that in each case it rapidly spreads. Forhtermore, up tp now the N501T mutation emerged and spread only from a single branch.

Interestingly, experimental measurements of binding affinity upon single mutations found that changing residue 501 from N to both T and Y results in an increase of the affinity, while most of the other possible mutations lead to its decrease [31].

In the following, we propose to investigate in greater detail what are the molecular features responsible for the increase/decrease of complex stability upon mutations.

### B. Fluctuation of interface residues for the different variants

In order to analyze the role of amino acid substitutions on the binding between the RBD of the SARS-CoV-2 spike protein and the ACE2 receptor, we performed extensive molecular dynamics simulations on a set of spike variants in complex with ACE2 and characterized the effects of the different mutations on the stability of the protein complex.

In particular, we considered the spike protein first detected in Wuhan at the beginning of the epidemics as the ‘wild type’ (WT). Starting from the structural complex comprising the RBD of the WT spike and the human ACE2 receptor (pdb id: 6m0j), we obtained the RBD of variant B.1.1.7 (the English variant) mutating residue 501 from amino acid N to Y.

In addition, we also considered four other single-mutated complexes: N501T and N501F, which are expected to display higher affinity; N501D and N501K, which should exhibit lower stability with respect to the wild type one [31].

Then we move to consider cases in which three mutations are present in the RBD, i.e. we analyze the South African and Amazonian variants carrying the mutations K417T-E484K-N501Y and K417N-E484K-N501Y, respectively.

All considered systems are reported in Table I.

**TABLE I:** Variants considered in the present work.

|  |  |
| --- | --- |
| Wild type (WT) | N501N |
| English (UK) | N501Y |
| SARS-03 like | N501T |
| Negative control 1 | N501K |
| Negative control 2 | N501D |
| Positive control 1 | N501F |
| South African | K417N - E484K - N501Y |
| Amazonian | K417T - E484K - N501Y |

For each considered system, mutations were obtained using Pymol (see Methods) and a 500 ns-long molecular dynamics simulation is conducted. To ensure a complete equilibration of the systems, only configurations after 250 ns are used in the analyses.

The first observable we consider to evaluate the stability of each RBD-ACE2 complex, was the Root Mean Square Fluctuation (RMSF). In Figure 2a, we show for each residue, both for spike protein and for the ACE2 receptor, the RMSF value obtained considering each molecular dynamics simulation at equilibrium. The average of the RMSF of all contact residues provides information on the overall mobility of the interface. The binding residues of ACE2 and spike have a comparable mean fluctuation value between them (of about 1.25 °*A*, as the average of all systems). The negative control 1 and 2 systems (whose mutations are N501K and N501D, respectively) are characterized by a higher *k*_*d*_ than the WT form of the spike. This is clearly evident in terms of atomic fluctuation, given that these systems show greater mobility of the interface residues, with an RMSF average of 1.53 °*A* and 1.31 °*A*, respectively. The system characterized only by N501Y mutation (UK variant) is the most stable of all, with an average of 1.01 °*A*. The Amazon variant and the South African variant also have a low average fluctuation compared to the other systems, of 1.14±0.23 °*A* and 1.22±0.20 °*A*, respectively. A more detailed analysis was performed on the atomic fluctuation of residue 501. In all systems, except for the South African variant, residue 501 of the spike protein has a lower average fluctuation than the average of all residues belonging to the same interface. In particular, the English variant, the Amazonian variant, and the positive control mutation (N501F) show a fluctuation of the residue 501 lower than the average of the RMSF values. Indeed, the RMSF value for residue 501 in these three systems is 1.00, 1.00, and 1.20, while the average values of RMSF for the three systems are 1.01±0.16, 1.14±0.23, and 1.25±0.20, respectively. Therefore, for these three systems, residue 501 is particularly stable with respect to the other residues. On the contrary, the other systems have a more pronounced fluctuation of residue 501. Among all, the two negative control systems, N501K and N501D mutations are characterized by a very high average RMSF value: 1.83 and 2.03 respectively. The results of the analysis of atomic fluctuation overtime of the interface residues highlight the stability of systems with higher binding affinity, including the South African and Amazonian variant for which we have not considered experimental data (since they are not present here [31]). In order to investigate the stability of intermolecular interactions during molecular dynamics, we calculated the contact frequency matrix for each system. Therefore, we define contact between two residues if their *α*-carbon atoms have a distance less than 9 °*A*. For each matrix element, we then report the contact frequency between each residue of the spike protein and each other of the ACE2 receptor, then subtracting (to facilitate the comparison) each of these matrices with that obtained for the WT system (see Methods section). As shown in Figure 2b, we notice similarities between some matrices. For example, the matrix relating to the N501Y system is particularly similar to the N501F system, which is characterized by the highest experimental binding affinity value. An analysis of the Pearson correlation between each pair of matrices allows us to quantify this evidence. In Table II the correlation values between all the contact matrices are shown. Interestingly, the positive control system (characterized by the N501F mutation) has a mean contact map highly correlated with that of the UK, Amazon, and South African variant, with a Pearson correlation value of 0.96, 0.92, and 0.85 respectively. The two systems formed by the two mutations with high *K*_*d*_ (the two negative controls), on the other hand, show lower correlation values with any other system, with an average of the Pearson coefficient of 0.34 and 0.32 respectively. These results represent the first level of classification of the mutated systems of this study and allow us to identify the properties of mean atomic fluctuation at a single residue level.

**TABLE II:** Correlation matrix between couples of variants. Correlation values (resp. p-values) are reported in the upper (resp. lower) triangular matrix.

|  | K417N - E484K - N501Y | K417T - E484K - N501Y | N501D | N501F | N501K | N501T | N501Y |
| --- | --- | --- | --- | --- | --- | --- | --- |
| K417N - E484K - N501Y | - | 0.91 | 0.44 | 0.92 | 0.38 | 0.67 | 0.77 |
| K417T - E484K - N501Y | $< 10^{-14}$ | - | 0.20 | 0.96 | 0.31 | 0.61 | 0.87 |
| N501D | $3.6 \times 10^{-14}$ | $8.1 \times 10^{-4}$ | - | 0.23 | 0.35 | 0.52 | 0.17 |
| N501F | $< 10^{-14}$ | $< 10^{-14}$ | $9.8 \times 10^{-5}$ | - | 0.34 | 0.67 | 0.85 |
| N501K | $8.7 \times 10^{-11}$ | $2.6 \times 10^{-7}$ | $4.6 \times 10^{-9}$ | $1.4 \times 10^{-8}$ | - | 0.28 | 0.26 |
| N501T | $< 10^{-14}$ | $< 10^{-14}$ | $< 10^{-14}$ | $< 10^{-14}$ | $2.4 \times 10^{-6}$ | - | 0.66 |
| N501Y | $< 10^{-14}$ | $< 10^{-14}$ | $5.1 \times 10^{-3}$ | $< 10^{-14}$ | $9.8 \times 10^{-6}$ | $< 10^{-14}$ | - |

**FIG. 2:**
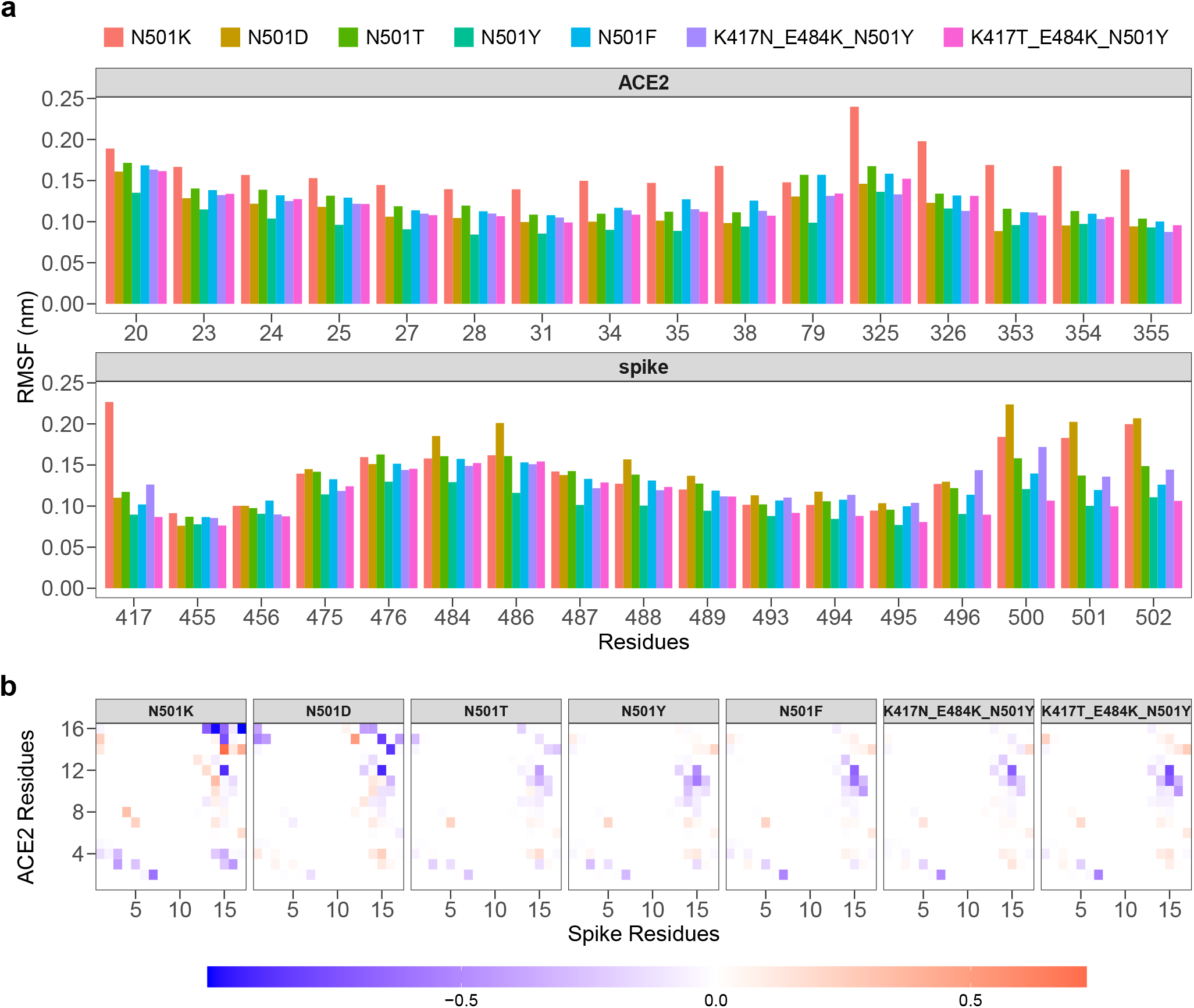
**a)** Root mean square fluctuations (RMSF) of ACE2 (top) and SARS-CoV-2 spike (bottom) protein residues found in interaction during the dynamics. Different colors corresponds to different spike variants (see Table I. **b)** Difference between the contact probability matrices of the interacting residues between each spike variant and the Wuhan (WT) one (see Table I). Residues are considered as interacting if the distance between their *α*-carbons is lower than 9°*A*.

### C. Principal Component Analysis of contact frequency

In order to obtain a more exhaustive overview of the binding between ACE2 and spike protein, we investigate the frequency of the contacts for each analyzed system. To this end, we associate each system to all intermolecular contact pairs, for each of which we consider the percentage of contacts during the simulation. Therefore, each system is described by a vector of 272 elements, since the residues of the binding site are 17 belonging to the spike protein and 16 belonging to the ACE2 receptor (see Methods). The first two principal components explain 54.4% and 35.3% of the total variance. The projection of each system on the essential plane of the first two principal components allows to clearly distinguish, completely unsupervised, the mutations with high affinity from those with low binding affinity. In particular, the projection along the first principal component shows a very interesting trend (as shown in Figure 3): the two mutations of the negative control have a positive value on the first component (distinguishing themselves from any other). On the other hand, the first three systems along this component are the Amazon variant, the English variant, and the positive control (N501F). We also analyze the loading of each interacting residue pair on the first principal component. In Figure 3 we show the considered pairs, given that their projection along the axis of the first component is high compared to the others (and the trend of the contact frequency is not trivial between the different systems). In most cases, the pairs of interacting residues at the interface show a trend (increasing or decreasing) of the frequency of the contacts for the different systems. For example, the percentage of contacts during the simulations of the pair composed of the S 494 residue of the spike protein and the H 34 residue of the ACE2 receptor (S494-H34), progressively decreases for systems with low binding affinity. Therefore, this pair is more stable in high binding affinity systems. With an opposite trend, the pair of residues G502-Q325 shows an increasing trend, meaning that this is more present for systems with low binding affinity. This analysis provides information on which pairs of intermolecular interactions are more stable in high-affinity binding systems and which pairs, on the other hand, are most present in low-affinity binding systems.

**FIG. 3:**
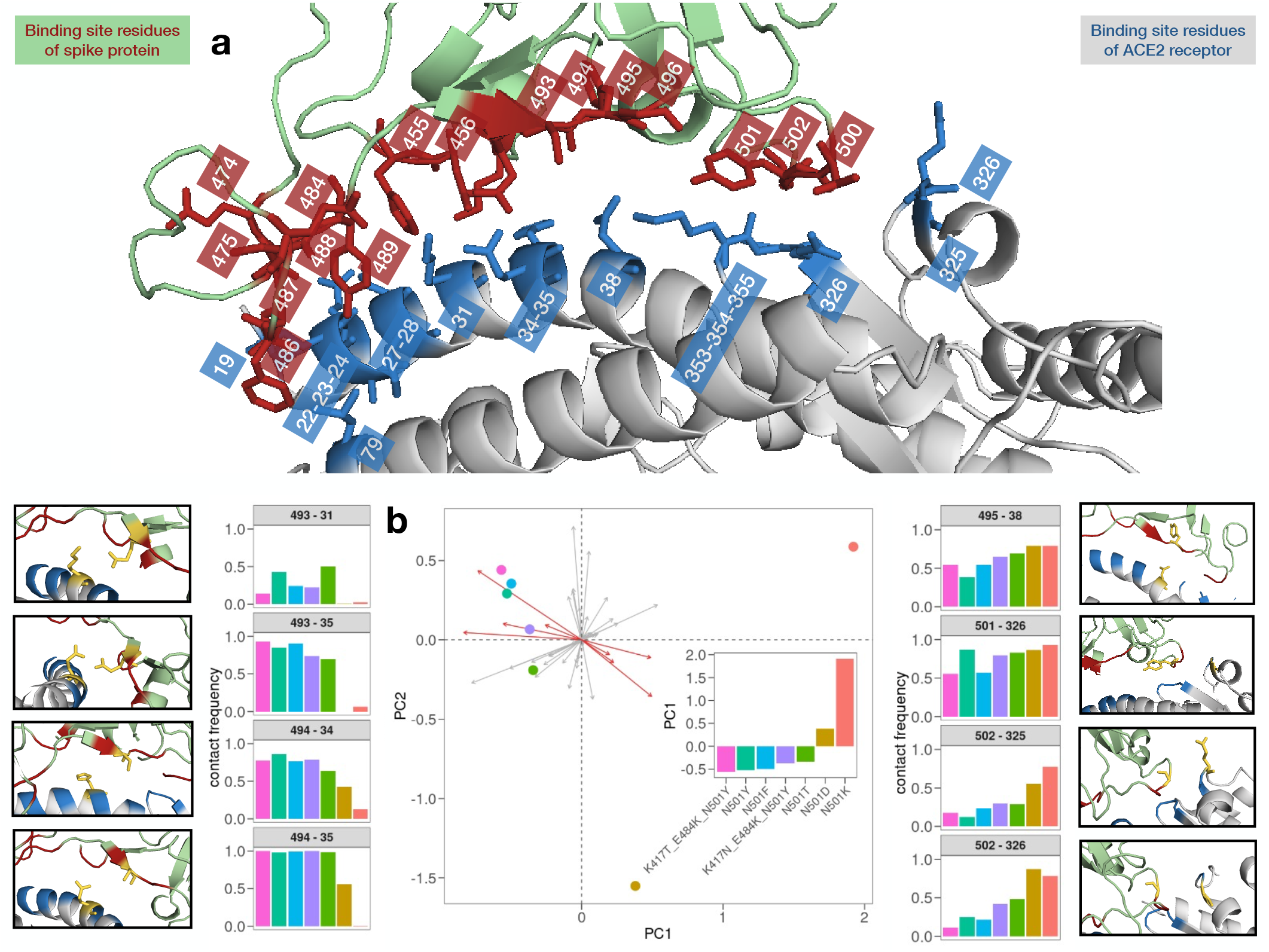
**a)** Ribbon representation of the binding site of the SARS-CoV-2 spike protein bound to human ACE2 (pdb id: 6m0j). Interacting residues, i.e. the ones whose *α*-carbons have a distance lower than 9 °*A*, are represented with sticks and listed on the left (spike’s) and right (ACE2’s) sides of the panel. **b)** Projections on the two principal components of the analyzed variants (colored dots). Principal component analysis was performed over the contact probabilities of each couple of interacting residues reported in panel a). The inset shows the projections on the first component. Side plots display the contact probability values for the eight couples of residues that contributed the most in differentiating the variants in the principal component analysis.

### D. Graph theory-based analysis of binding site residues

A higher level of complexity of the binding properties was addressed by analyzing the organization of interactions between the interface residues. To this end, we model the interaction between the two proteins as a bi-partite network, schematizing each residue as a node of the network and each intermolecular interaction as an edge. Only intermolecular interactions were considered for this analysis, so interactions between two residues belonging to the same protein are not included in the graph definition. In particular, we define a weighted graph for each system, weighing each link connecting two residues with the corresponding contact frequency calculated from the molecular dynamics, defining, in this case, contact between two residues if their distance is less than 8.5°*A* in agreement with [32]. As an example for a better interpretation of this modeling for the molecular complex, we report the distances that every residue belonging to the ACE2 binding site has with the residue 501 of the Spike protein since this is of primary importance in the variants considered in this work (see Figure 4). The mean distance between residue 501 of the spike protein and any other (within 12 °*A*) of the ACE2 receptor was calculated. This analysis shows that the systems with lower experimental binding affinity are typically characterized by a greater distance (N501K and N501D single mutation) between the residues belonging to the binding site of ACE2 and the residue 501 of the spike protein. Interestingly, the system with the N501D mutation has a shorter distance than any other system between residues 501 and residues 37, 38, 39, 41, 42, and 45 of the receptor. Similarly, for the system having the N501K mutation, it shows the lowest distance between residue 501 and residue 383, 386, and 387 belonging to the ACE2 receptor. Extending the analysis to any other residue of the spike protein interface, we investigate the local organization of the inter-molecular contacts through centrality measures. To this end, both the *betweenness centrality* and *closeness centrality* parameters were considered as a local descriptor of the interaction of each residue, since these are certainly two of the most widely used descriptors for the centrality analysis of a node.

**FIG. 4:**
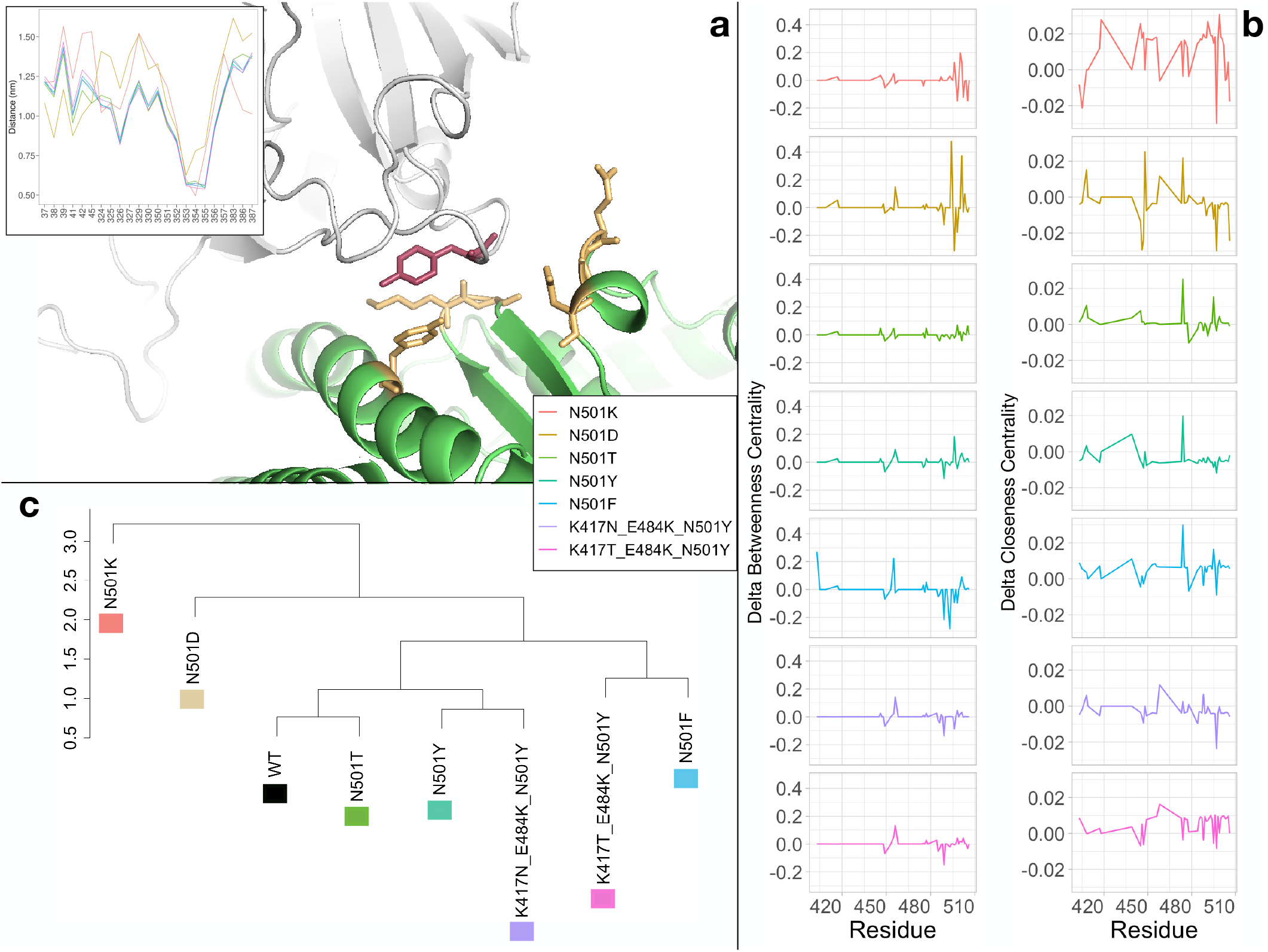
**a)** Cartoon representation of the complex (pdb id:6m0j) between SARS-CoV-2 spike protein (grey) and the human ACE2 receptor (green). Residue 501 of the spike protein is represented in red sticks, while a set of interacting residues of the ACE protein are identified in yellow sticks on the basis of a spatial distance threshold of 9 *Å*. The inset shows the average minimum distance of the ACE2 interacting residues with residue 501 o the spike protein for each variant. **b)** Difference in Betweenness (left column) and Closeness (right column) centrality of each node of the SARS-CoV-2 spike found in interaction with ACE2 receptor between the seven studied spike variants and the reference one (see Table I). **c)** Hierarchical clustering of the seven spike variant together with the reference one (see Table I).

In order to compare the different interaction organizations of each molecular system, we performed a clustering analysis considering a single vector for each system, which is composed of the combination of the two descriptors considered. The profile is relative only to the residues belonging to the spike interface (see fig 3a).

Also in this analysis, we find a clear separation of mutations with lower binding affinity from the others, showing the ability of our approach based on molecular dynamics to distinguish high and low-affinity systems. The WT form of the complex takes place in an intermediate position between low-affinity and high-affinity mutations. In particular, the WT system is characterized by a profile very similar to the N501T mutation, which has the same amino acid in position 501 of the SARS-CoV system.

On the other side, the Amazon variant and the South African variant appear to have a very similar organization of contacts with positive control and English variant, respectively. Interesting to note that the Amazon variant, which is causing an important concern in the world, is very similar to the system characterized by the single mutation N501F, the one with the best experimental binding affinity.

### E. Shape variation of binding site region

Finally, we focused on the whole binding region of both SARS-CoV-2 spike protein and ACE2 receptor and assessed the shape complementarity of the binding site. To do so, we first expanded the molecular surfaces of the two molecular partners (see Figure 5a) on the basis of the 3D Zernike moments (see Methods) and then computed the Euclidean distance between the two sets of invariant descriptors. We repeated the procedure for 250 configurations sampled from the equilibrium of each of the molecular dynamics simulations we performed. We thus obtained a distribution of Zernike distances, i.e. of complementarity scores, for each investigated variant. As one could expect, all distributions are centered around similar values of complementarity since the overall shape of the binding site does not vary so much performing a few point mutations. However, looking at the variance of the distributions for the five single-mutation variants, we found a similar trend between the experimentally measured complex affinities and the variance of the shape complementarity as one can see from Figure 5b where we compared the difference in affinity and shape variance of the five investigated variants with respect to the wild type. It is interesting to note that the N501Y variant displays a lower shape variance with respect to the wild type than the N501F variant. Unfortunately, as far as our knowledge goes, no binding affinity data are available for the three-mutation variants (i.e. the South African and Amazonian variants). We thus proceeded to compute the shape variance from the molecular dynamics configurations and compare them with single-mutation ones. Interestingly, we found that both variants show a lower variance with respect to the negative controls (N501D and N501K), with the South African variant being more motile than the English one while the Amazonian variant having the smallest variance (see Figure 5c). This result provides an important example of the not-trivial effect of cooperativity, in fact, the same mutation (Y501F) gives an opposite outcome in the two variants, which now differ in terms of the interactions between a couple of residues 417T-501F against 417N-501F. Finally, we run two additional molecular dynamics simulations substituting amino acid F to Y in the two triple-mutated variants in order to check whether such mutation could bring to an enhancement in the binding propensity. Looking again at the shape variance, we observed that the modified South African and Amazonian variants behave oppositely: while a Y to F mutation on the 501 residue increases the complementarity variability for the Amazonian variant thus worsening its binding capacity, the same mutation on the South African one brings a stabilization effect, with the complex maintaining a more stable shape complementarity during the dynamics.

**FIG. 5:**
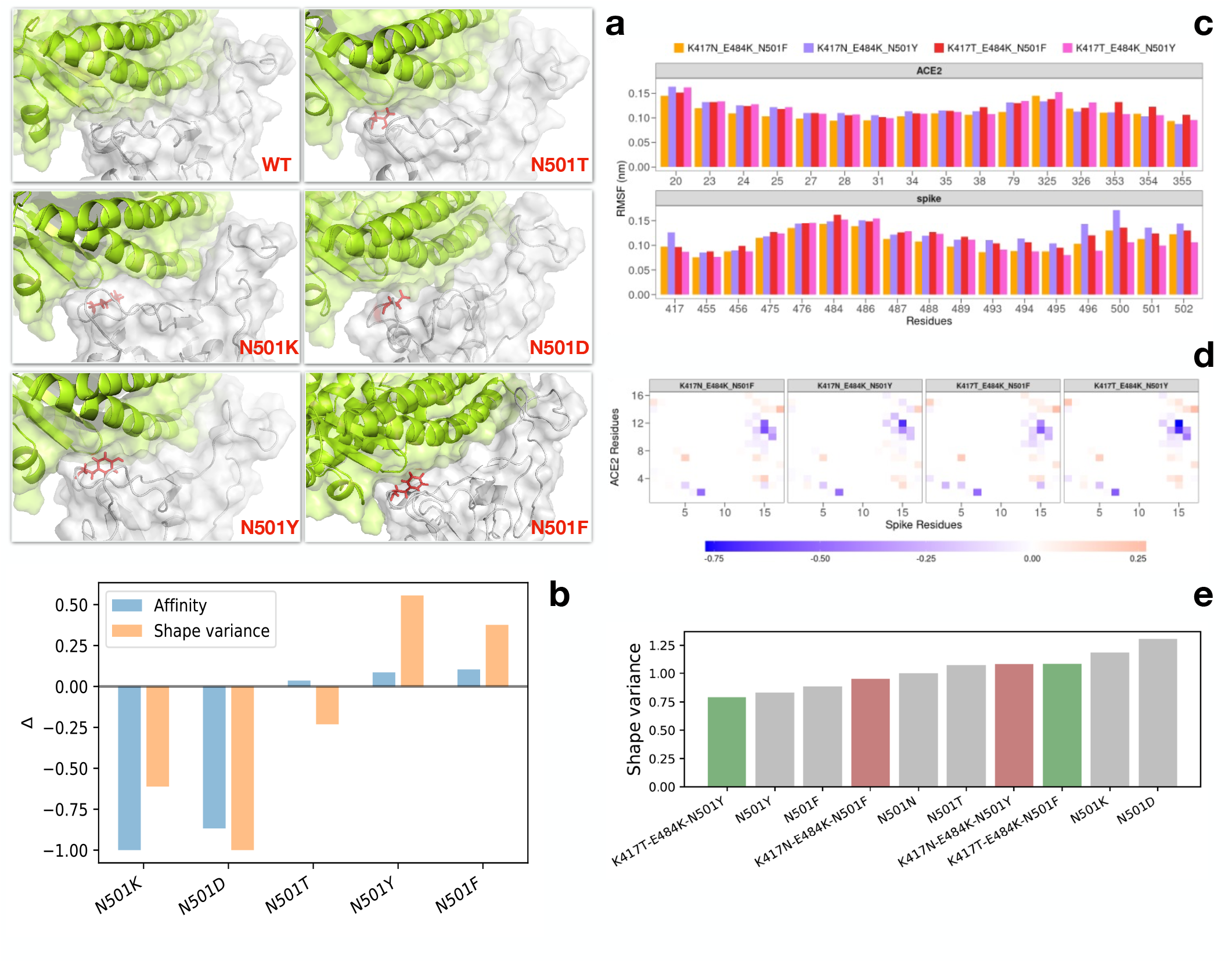
**a)** Cartoon representations of the binding region of the complex formed by SARS-CoV-2 spike protein (grey) and human ACE2 receptor (green). for the wild-type complex and the five single-mutation variants. The spike residue 501 is represented with red sticks. The molecular surfaces for both spike and ACE2 are showed. **b)** Experimental affinity values (taken from [31]) for the spike-ACE2 complexes (blue bars) and shape variance as measured by the Zernike descriptors on molecular dynamics configurations (orange bars). Both quantities are obtained by difference with respect to the reference complex data (see Table I). **c**) Root mean square fluctuations (RMSF) of ACE2 (top) and SARS-CoV-2 spike (bottom) protein residues found in interaction during the dynamics for the South African and Amazonian variants together with the two versions carrying phenylalanine at position 501. **d)** Difference between the contact probability matrices of the interacting residues between each three-mutation spike variant and the Wuhan (WT) one (see Table I). Residues are considered as interacting if the distance between their *α*-carbons is lower than 9°*A*. **e)** Shape variance as measured by the Zernike descriptors on molecular dynamics configurations for all the variants reported in Table I.

## III. CONCLUSIONS

The major goals of a virus are replication and spread, and SARS-CoV-2 coronavirus is constituting no exception. The vast, worldwide, diffusion of the SARS-CoV-2 epidemics indeed is providing the virus the possibility of exploring the genomic landscape and accumulating mutations. The resulting viral variants are subject to natural selection so that mutations that increase the diffusion rapidly get fixed in the viral genomic pool. Besides the transmission of the virus itself, antiviral treatments also contribute to selection. Indeed with the introduction of vaccines, new mutations inducing escape and resistance to resistant to available treatments producing are expected to couple with more virulent strains [33, 34].

The new frontier for fighting the COVID-19 pandemic seems to become even more based on controlling and understanding the accumulation of variations in the SARS-CoV-2 genome.

To date, several different variants have already emerged.

In particular, those with mutations on the spike protein are attracting a lot of attention as this protein is responsible for the binding to cellular receptor and attachment factors but also the primary target of the antibodies of the immune system.

Here, we deployed extensive molecular dynamics simulations to characterize the effect of different possible mutations of the spike residues 417, 484, and 501, which are the mutations found in the most relevant observed variants. In particular, mutation N501Y has been found in both the English, South African, and Amazonian variants. Analyzing the equilibrium configurations acquired by the spike-ACE2 complex in terms of residue fluctuations, networks of contacts, and conservation of shape complementary of the binding region, we found that indeed mutations on residue 501 strongly influence the dynamical stability of the complex. Most importantly, a phenylalanine substitution in position 501 increases the stability of the South African variant via a cooperative action with residue 417T. In conclusion, our results suggest close surveillance for the emergence of such mutation to anticipate (and minimize) its effects in the viral spread and eventual early incorporation of this information into diagnostic and pharmacological procedures.

## IV. MATERIALS AND METHODS

### A. Complex structural data

The complex of SARS-CoV-2 spike protein bound to human ACE receptor has been taken from the PDB bank (pdb id: 6m0j). In particular, only the receptor-binding domain (RBD) of the spike and the extracellular domain of ACE2 are considered. Each variant considered in the present study has been obtained manually mutating the experimental complex via the Pymol software [35]. All information about amino acid substitutions is reported in Table I.

### B. Molecular dynamics simulations

All simulations were performed using Gromacs [36]. Topologies of the system were built using the CHARMM-27 force field [37]. The protein was placed in a dodecahedric simulative box, with periodic boundary conditions, filled with TIP3P water molecules [38]. For all simulated systems, we checked that each atom of the proteins was at least at a distance of 1.1 nm from the box borders. Each system was then minimized with the steepest descent algorithm. Next, a relaxation of water molecules and thermalization of the system was run in NVT and NPT environments each for 0.1 ns at 2 fs time-step. The temperature was kept constant at 300 K with v-rescale thermostat [39]; the final pressure was fixed at 1 bar with the Parrinello-Rahman barostat [40].

LINCS algorithm [41] was used to constraint bonds involving hydrogen atoms. A cut-off of 12 °A was imposed for the evaluation of short-range non-bonded interactions and the Particle Mesh Ewald method [42] for the long-range electrostatic interactions. The described procedure was used for all the performed simulations.

### C. Interface residue definition and contact probability calculation

For each analyzed complex, the interface was defined by taking ACE2 and SARS-CoV-2 spike residues whose *α*-carbons have a distance lower than 12 °A at time *t* = 250 ns, that is, after the equilibration phase. To end with a comparable set for all complexes, we selected the residues common to all interfaces obtaining 16 residuals for ACE2 and 17 for the spike protein. For each couple of interacting residues among the two proteins, we calculated the contact frequency, counting how many times each couple of residues had a distance lower than 9 °A between two *α*-carbons in each frame of the dynamics at equilibrium.

We got a 16×17 matrix for every complex reported in Table I. Contact matrices shown in Figure 2b are obtained by subtracting the wild type one.

### D. Principal component analysis and clustering

From the contact matrices of all complexes, we performed a principal component analysis (PCA) in which the starting matrix consisted of 7 rows (the variants) and 272 columns (the contact frequencies of each pair of spike-ACE2 *α*-carbons). The clustering analysis was performed using the ‘hclust’ function of R, preserving the default clustering algorithm (the “complete” method). We first computed the contact percentage matrix for each system, as obtained from the molecular dynamics frames. For graphics analysis, we define contact between two residues if the distance between their center is less than 8.5 °*A*, as proposed in [32]. Then, we defined a weighted graph for each matrix and calculated the betweenness centrality and closeness centrality parameters for each residue.

In particular, the betweenness centrality of a node *i* is given by:

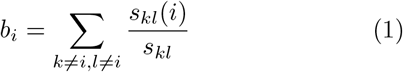

where *s*_*kl*_ (*i*) is number of weighted shortest paths (*s*) that go from node *k* to node *l* passing through node *i*; while *s*_*kl*_ is the total number of weighted shortest paths from node *k* to node *l*.

Similarly, the closeness centrality of a node *i* is defined by the inverse of the average length (*<* · *>*) of the weighted shortest paths to/from all the other nodes in the network:

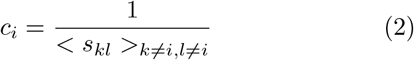

Both quantities were obtained via the corresponding functions of the *‘igraph’* package of R [43].

For each system, we combined the two vectors (which have values of betweenness and closeness for single residues) normalized to 1. We used the Euclidean distance to compare every pair of vectors. Analyses were performed using R standard libraries [44].

#### Molecular surface analysis via Zernike descriptors

Given an ACE2-RBD complex simulated in molecular dynamics, for each protein, we calculate separately the molecular surface using DMS software[45]. Once extracted the portion of protein surface in interaction, with a voxelization procedure we represent the protein patch as a 3D function.

This 3D function can be described as a series expansion on the basis of the 3D Zernike Polynomials[27, 46, 47]. Taking the norm of the expansion coefficients we deal with an ordered set of numerical descriptors that compactly summarize the shape of the examined molecular surface.

Indeed, a function *f* (*r, θ, φ*) can be written as:

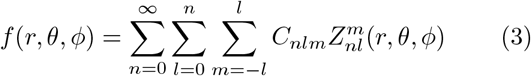

where 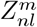 are the 3D Zernike polynomials, while the coefficients *C*_*nlm*_ are called Zernike moments.

The precision of the description can be selected by modifying the order of the expansion N. In this work, we fix N = 20, corresponding to 121 numerical descriptors representing each function.

The 3D Zernike Moments can be seen as:

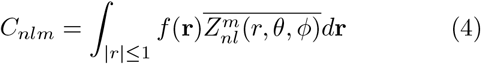

where 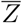 represent the complex conjugate.

The norms of such moments, with respect to the index m, are invariant under translation and rotation. Indeed, the Zernike Descriptors are defined as:

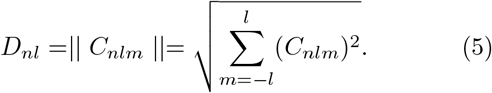

The shape complementarity between 2 surfaces can be easily evaluated applying a metric between the two vectors of numbers describing them[48–50]. Indeed, we adopted the euclidean distance. When 2 surfaces have a low distance between them, they are characterized by a similar shape and therefore they are suitable for binding.

## Author Contributions

All authors analyzed results; all authors wrote and revised the paper.

## References

[1] C. Huang, Y. Wang, X. Li, L. Ren, J. Zhao, Y. Hu, L. Zhang, G. Fan, J. Xu, X. Gu, et al., The Lancet 395, 497 (2020).

[2] N. Zhu, D. Zhang, W. Wang, X. Li, B. Yang, J. Song, X. Zhao, B. Huang, W. Shi, R. Lu, et al., New England Journal of Medicine (2020).

[3] A. Fontanet, B. Autran, B. Lina, M. P. Kieny, S. S. A. Karim, and D. Sridhar, The Lancet 397, 952 (2021).

[4] S. Duchene, L. Featherstone, M. Haritopoulou-Sinanidou, A. Rambaut, P. Lemey, and G. Baele, Virus evolution 6, veaa061 (2020).

[5] S. Portelli, M. Olshansky, C. H. Rodrigues, E. N. D’Souza, Y. Myung, M. Silk, A. Alavi, D. E. Pires, and D. B. Ascher, Nature Genetics 52, 999 (2020).

[6] J. A. Plante, Y. Liu, J. Liu, H. Xia, B. A. Johnson, K. G. Lokugamage, X. Zhang, A. E. Muruato, J. Zou, C. R. Fontes-Garfias, et al., Nature pp. 1–6 (2020).

[7] J. Zhang, Y. Cai, T. Xiao, J. Lu, H. Peng, S. M. Sterling, R. M. Walsh, S. Rits-Volloch, H. Zhu, A. N. Woosley, et al., Science (2021).

[8] E. Trucchi, P. Gratton, F. Mafessoni, S. Motta, F. Cicconardi, F. Mancia, G. Bertorelle, I. D’Annessa, and D. D. Marino, Molecular biology and evolution (2021).

[9] R. S. Baric, New England Journal of Medicine 383, 2684 (2020).

[10] E. Trucchi, P. Gratton, F. Mafessoni, S. Motta, F. Cicconardi, F. Mancia, G. Bertorelle, I. D’Annessa, and D. D. Marino, Molecular Biology and Evolution (2021).

[11] J. W. Tang, P. A. Tambyah, and D. S. Hui, The Journal of infection (2020).

[12] S. E. Galloway, P. Paul, D. R. MacCannell, M. A. Johansson, J. T. Brooks, A. MacNeil, R. B. Slayton, S. Tong, B. J. Silk, G. L. Armstrong, et al., Morbidity and Mortality Weekly Report 70, 95 (2021).

[13] E. Volz, S. Mishra, M. Chand, J. C. Barrett, R. Johnson, L. Geidelberg, W. R. Hinsley, D. J. Laydon, G. Dabrera, Á. O’Toole, et al., medRxiv pp. 2020–12 (2021).

[14] R. Challen, E. Brooks-Pollock, J. M. Read, L. Dyson, K. Tsaneva-Atanasova, and L. Danon, bmj 372 (2021).

[15] M. Chand et al., Public Health England. PHE (2020).

[16] H. Tegally, E. Wilkinson, M. Giovanetti, A. Iranzadeh, V. Fonseca, J. Giandhari, D. Doolabh, S. Pillay, E. J. San, N. Msomi, et al., medRxiv (2020).

[17] F. Naveca, C. da Costa, V. Nascimento, V. Souza, A. Corado, F. Nascimento, Á. Costa, D. Duarte, G. Silva, M. Mejía, et al., virological. org. Preprint available at: https://virological.org/t/sars-cov-2-reinfection-by-thenew-variant-of-concern-voc-p-1-in-amazonas-brazil/596. Available at: https://virological.org/t/sars-cov-2-reinfection-by-the-new-variant-of-concern-voc-p-1-in-amazonas-brazil/596 (2021).

[18] R. J. Hulswit, Y. Lang, M. J. Bakkers, W. Li, Z. Li, A. Schouten, B. Ophorst, F. J. van Kuppeveld, G.-J. Boons, B.-J. Bosch, et al., Proceedings of the National Academy of Sciences 116, 2681 (2019).

[19] C. Schwegmann-Weßels and G. Herrler, Glycoconjugate journal 23, 51 (2006).

[20] E. Milanetti, M. Miotto, L. Di Rienzo, M. Monti, G. Gosti, and G. Ruocco, arXiv preprint 2003.11107 (2020).

[21] A. N. Baker, S.-J. Richards, C. S. Guy, T. R. Congdon, M. Hasan, A. J. Zwetsloot, A. Gallo, J. R. Lewandowski, P. J. Stansfeld, A. Straube, et al., ACS Central Science (2020).

[22] D. Schoeman and B. C. Fielding, Virology journal 16, 1 (2019).

[23] Z. Zhu, X. Lian, X. Su, W. Wu, G. A. Marraro, and Y. Zeng, Respiratory research 21, 1 (2020).

[24] X. Ou, Y. Liu, X. Lei, P. Li, D. Mi, L. Ren, L. Guo, R. Guo, T. Chen, J. Hu, et al., Nature communications 11, 1 (2020).

[25] R. Yan, Y. Zhang, Y. Li, L. Xia, Y. Guo, and Q. Zhou, Science (2020).

[26] P. Zhou, X.-L. Yang, X.-G. Wang, B. Hu, L. Zhang, W. Zhang, H.-R. Si, Y. Zhu, B. Li, C.-L. Huang, et al., Nature pp. 1–4 (2020).

[27] L. Di Rienzo, E. Milanetti, R. Lepore, P. P. Olimpieri, and A. Tramontano, Scientific reports 7, 1 (2017).

[28] A. Ali and R. Vijayan, Scientific reports 10, 1 (2020).

[29] B. Korber, W. M. Fischer, S. Gnanakaran, H. Yoon, J. Theiler, W. Abfalterer, N. Hengartner, E. E. Giorgi, T. Bhattacharya, B. Foley, et al., Cell 182, 812 (2020), ISSN 00928674, URL https://doi.org/10.1016/j.cell.2020.06.043 https://linkinghub.elsevier.com/retrieve/pii/S0092867420308205.

[30] J. Hadfield, C. Megill, S. M. Bell, J. Huddleston, B. Potter, C. Callender, P. Sagulenko, T. Bedford, and R. A. Neher, Bioinformatics 34, 4121 (2018).

[31] T. N. Starr, A. J. Greaney, S. K. Hilton, D. Ellis, K. H. Crawford, A. S. Dingens, M. J. Navarro, J. E. Bowen, M. A. Tortorici, A. C. Walls, et al., Cell 182, 1295 (2020).

[32] B. Chakrabarty and N. Parekh, Nucleic acids research 44, W375 (2016).

[33] R. Sanjuán and P. Domingo-Calap, Cellular and Molecular Life Sciences 73, 4433 (2016).

[34] M. Miotto and L. Monacelli, Phys. Rev. Research 2, 043026 (2020).

[35] Schrödinger, LLC (2015).

[36] D. V. D. Spoel, E. Lindahl, B. Hess, G. Groenhof, A. E. Mark, and H. J. C. Berendsen, Journal of Computational Chemistry 26, 1701 (2005).

[37] B. R. Brooks, C. L. Brooks, A. D. Mackerell, L. Nilsson, R. J. Petrella, B. Roux, Y. Won, G. Archontis, C. Bartels, S. Boresch, et al., Journal of Computational Chemistry 30, 1545 (2009).

[38] W. L. Jorgensen, J. Chandrasekhar, J. D. Madura, R. W. Impey, and M. L. Klein, The Journal of Chemical Physics 79, 926 (1983).

[39] G. Bussi, D. Donadio, and M. Parrinello, The Journal of Chemical Physics 126, 014101 (2007).

[40] M. Parrinello and A. Rahman, Physical Review Letters 45, 1196 (1980).

[41] B. Hess, H. Bekker, H. J. C. Berendsen, and J. G. E. M. Fraaije, Journal of Computational Chemistry 18, 1463 (1997).

[42] T. E. I. Cheatham, J. L. Miller, T. Fox, T. A. Darden, and P. A. Kollman, Journal of the American Chemical Society 117, 4193 (1995).

[43] G. Csardi, T. Nepusz, et al., InterJournal, complex systems 1695, 1 (2006).

[44] R Core Team, R: A Language and Environment for Statistical Computing, R Foundation for Statistical Computing, Vienna, Austria (2020), URL https://www.R-project.org/.

[45] F. M. Richards, Annual review of biophysics and bioengineering 6, 151 (1977).

[46] V. Venkatraman, L. Sael, and D. Kihara, Cell biochemistry and biophysics 54, 23 (2009).

[47] M. Novotni and R. Klein, Computer-Aided Design 36, 1047 (2004).

[48] V. Venkatraman, Y. D. Yang, L. Sael, and D. Kihara, BMC bioinformatics 10, 407 (2009).

[49] S. Daberdaku and C. Ferrari, BMC bioinformatics 19, 35 (2018).

[50] L. Di Rienzo, E. Milanetti, J. Alba, and M. D’Abramo, Journal of Chemical Information and Modeling 60, 1390 (2020).

